# Abiotic factors modulate interspecies competition mediated by the type VI secretion system effectors in *Vibrio cholerae*

**DOI:** 10.1101/2021.05.27.445954

**Authors:** Ming-Xuan Tang, Tong-Tong Pei, Zeng-Hang Wang, Han Luo, Xing-Yu Wang, Tao G. Dong

## Abstract

*Vibrio cholerae*, the etiological pathogen of cholera, relies on its type VI secretion system (T6SS) as an effective weapon to survive in highly competitive communities. The anti-bacterial and anti-eukaryotic functions of T6SS depend on its secreted effectors that target multiple essential cellular processes. However, the mechanisms that account for effector diversity and different effectiveness during interspecies competition remain elusive. Here, we report that environmental cations and temperature play a key role in dictating effector-mediated competition of *Vibrio cholerae*. We found that *V. cholerae* could employ its cell-wall-targeting effector TseH to outcompete the otherwise resistant *Escherichia coli* and the *V. cholerae* immunity deletion mutant *ΔtsiH* when Ca^2+^ and Mg^2+^ were supplemented. The *E. coli* Δ*phoQ* mutant was more sensitive to TseH-mediated killing during competition, suggesting the metal-sensing PhoPQ two-component system is protective to *E. coli* from TseH activity. Using transcriptome analysis, we found multiple stress response systems, including acid stress response, oxidative stress response, and osmotic stress response, were activated in *E. coli* expressing TseH in comparison with *E. coli* expressing the inactive mutant TseH^H64A^. The membrane-targeting lipase effector TseL also exhibited reduced killing against *E. coli* when divalent cations were removed. In addition, competition analysis of *E. coli* with *V. cholerae* single-effector active strains reveals a temperature-dependent susceptibility of *E. coli* to effectors, VasX, VgrG3, and TseL. These findings suggest that abiotic factors, that *V. cholerae* frequently encounters in natural habitats, play a crucial role in dictating the competitive fitness conferred by the type VI secretion system in complex multispecies communities.

## Introduction

Living in complex natural and host environments, microbes frequently compete for limited nutrients and ecological niches. To survive, microbes have evolved multiple effective weapons [1–4], one of which is the type VI protein secretion system (T6SS) commonly found in gram-negative bacteria [5–7]. The T6SS consists of three structural parts, a bacteriophage-like baseplate (TssEFGK), a transmembrane complex (TssJLM), and a contractile-tube structure with the Hcp inner tube surrounded by VipA/VipB outer sheath [8–11]. The top of the T6SS Hcp tube is sharpened by a VgrG trimer-PAAR spike complex [12–14]. Upon sheath contraction, the T6SS could inject the inner tube-spike components and the associated effectors directly into the recipient cells [15–17].

The physical puncture of T6SS injection causes little harm to recipient cells and it is the T6SS effectors that mainly dictate the T6SS function [17–20]. Effectors are T6SS-secreted toxins exhibiting anti-bacterial and/or anti-eukaryotic activities, and the anti-bacterial effectors mainly include cell-wall disrupting effectors, membrane targeting lipases and pore-forming toxins, nucleases, and cytosolic toxins including NAD(P)^+^ hydrolase [21], ADP-ribosyl transferase [22], (p)ppApp synthetase [23]. To confer self-protection against anti-bacterial toxins, bacteria encode effector-cognate immunity proteins that specifically interact with effectors [24–27]. In contrast to the specific immunity protein-mediated protection, the innate-immunity-like stress response pathways also provide critical protection against effectors and T6SS delivery, including production of extracellular polysaccharides [18, 28], oxidative stress response [29, 30], envelope stress responses [18], acid and osmotic stress response [19]. In addition, spatial separation at the community level can also protect susceptible cells from T6SS aggressors [31–35].

T6SS species often encode multiple effector modules which may be used synergistically toward certain species or independently for killing of a broad spectrum of competitors [36–39]. However, some effectors are inactive under lab conditions. For example, in the cholera-causing pathogen *Vibrio cholerae*, the cell-wall targeting trans-peptidase TseH shows little toxicity against *Escherichia coli* or the *V. cholerae* immunity gene deletion mutant Δ*tsiH* when it is delivered by the T6SS [18, 40]. However, TseH is highly toxic to *Aeromonas* and other waterborne species that *V. cholerae* may encounter in the natural environments [18]. Similarly, the lipase effector TseL of *V. cholerae* could be secreted by the T6SS to kill the immunity gene mutant Δ*tsiV1* but it is not very effective against the *E. coli* prey [17, 19, 27]. The difference in susceptibility has been attributed to the nonspecific stress response mechanisms in diverse bacteria [18–20].

In this study, we report that a number of abiotic environmental factors, including cations, temperatures and antibiotics, also play a crucial role in T6SS-mediated interspecies competition by modulating target-species sensitivity to specific effectors. Using a panel of single-effector active *V. cholerae* V52 mutants [17, 18], we show that trace amounts of divalent cations, Mg^2+^ and Ca^2+^, could stimulate TseH toxicity against *E. coli* when delivered by the T6SS, and EDTA treatment abolished such stimulation. The *V. cholerae ΔtsiH* mutant was also more sensitive to TseH during competition when Ca^2+^ was supplemented. We further show that deletion of *phoQ*, encoding the sensor of the key two-component system PhoPQ that senses low Mg^2+^ [41], renders *E. coli* more sensitive to TseH-mediated killing, while a constitutively active PhoQ^D179L^ is more resistant. Transcriptome analysis of *E. coli* expressing TseH or its catalytic mutant TseH^H64A^ in the periplasm shows a global cellular change involving multiple pathways, including acid stress response, oxidative stress response, and osmotic stress response, which might be crucial for survival against TseH toxicity. The toxicity of TseL, a lipase effector in *V. cholerae*, was also modulated by divalent cations since EDTA treatment resulted in significantly reduced *E. coli* killing mediated by TseL. In addition to cations, *V. cholerae* effectors, VasX, VgrG3 and TseL, also show temperature-dependent toxicity when delivered individually to *E. coli*. Collectively, our data indicate effector-mediated toxicities are modulated by these abiotic factors, highlighting the complex effects of natural environments on interspecies interactions mediated by the T6SS.

## Results

### T6SS-delivered TseH is conditionally toxic to *E. coli* depending on agar sources

We have previously reported that the *V. cholerae tseH*^*+*^ only mutant, lacking all the other antibacterial effectors, could not outcompete *E. coli* due to the non-specific protections conferred by the envelope stress response pathways [18]. However, we serendipitously found that TseH toxicity was largely affected by the source of agars used for competition. When agar source 1 was used, we reproduced the earlier observation and showed that survival of *E. coli* competed with the *tseH*^*+*^ only mutant was comparable to that competed with the T6SS-null mutant Δ*vasK* and the four-effector inactive mutant *4eff*_*c*_ (Figure 1A, Supplementary Figure 1A). However, when a different source of agar was used (source 2), we noticed that *E. coli* survival was reduced by 10,000-fold in competition with the *V. cholerae tseH*^*+*^ only mutant, relative to that competed with the *4eff*_*c*_ and Δ*vasK* mutant. Survival of *E. coli* remained the same under both agar conditions for samples competed with wild-type *V. cholerae*, the Δ*vasK*, or the *4eff*_*c*_ mutant, separately.

**Figure 1.**
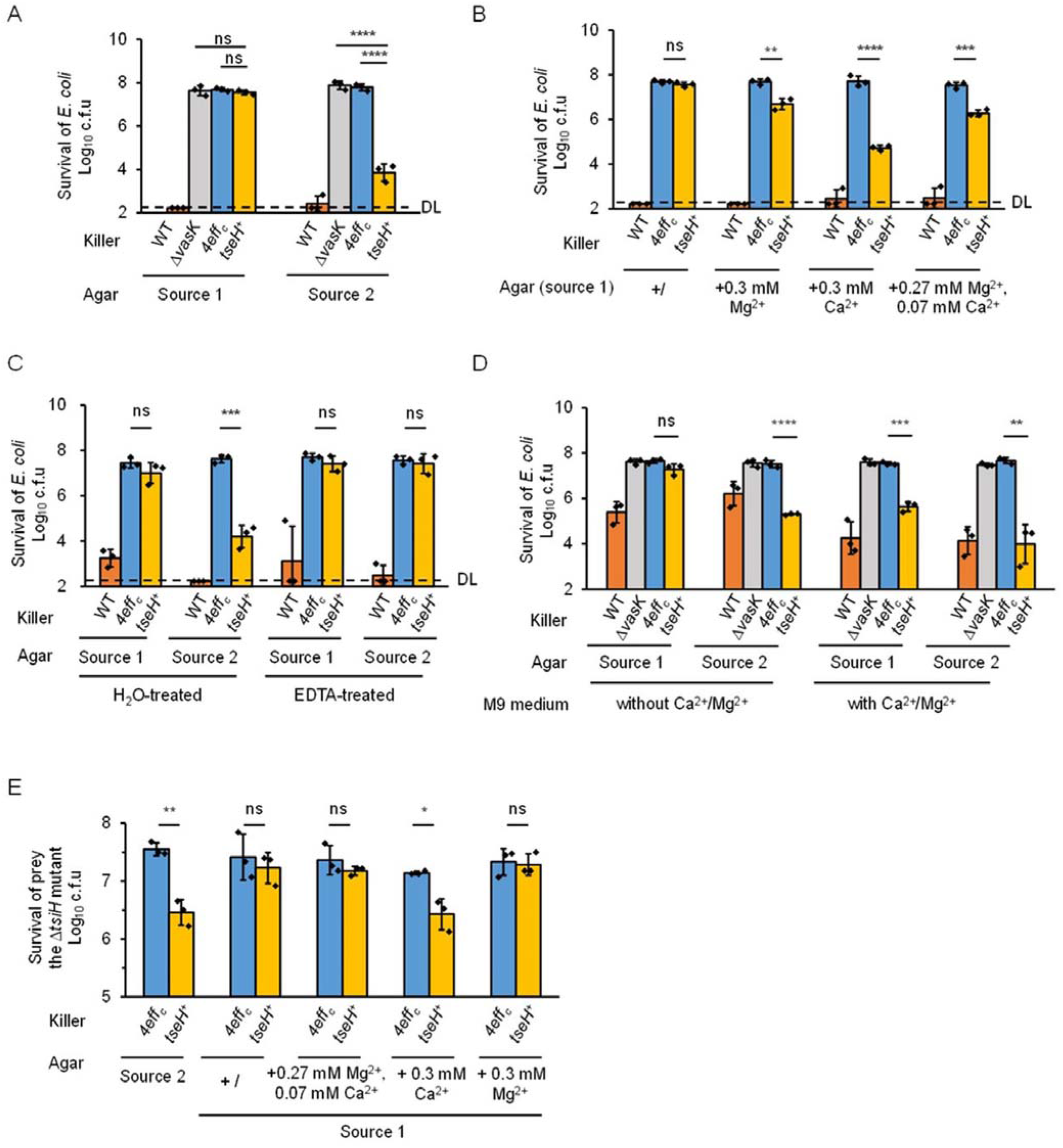
Mg^2+^ and Ca^2+^ stimulate TseH-mediated killing against *E. coli* and the *V. cholerae* Δ*tsiH* mutant. **A**, Survival of *E. coli* after competition with *V. cholerae* strains on LB plates of two different agar sources. **B**, Survival of *E. coli* competed with *V. cholerae* strains on LB plates of agar source 1, supplemented with Mg^2+^ and Ca^2+^. **C**, Effect of EDTA-treatment on survival of *E. coli* competed with *V. cholerae* strains. LB plates of ddH_2_O-treated agar were used as control. **D**, Survival of *E. coli* competed with *V. cholerae* strains on M9 source-1-agar or source-2-agar plates. **E**, Survival of the *V. cholerae* Δ*tsiH* mutant after competition with the *4eff*_*c*_ and *tseH*^*+*^ strain. For **A** to **E**, killer strains are indicated at the bottom of each panel. WT, wild type; Δ*vasK*, the T6SS-null Δ*vasK* mutant; *4eff*_*c*,_ the 4-antibacterial-effector-inactive mutant; *tseH*^*+*^, the TseH-active mutant. Survival of prey cells was enumerated by serial plating on selective medium. Killer survival was shown in **Supplementary Figure 1A-1E** respectively. Error bars indicate the mean ± standard deviation of three biological replicates. Statistical significance was calculated using a two-tailed Student’s *t*-test for two groups comparison or one-way ANOVA test for more than two groups comparison. ns, not significant; *, *p* <0.05; **, *p* <0.01; ***, *p*< 0.001; ****, *p*< 0.0001. DL, detection limit.

### Mg^2+^ and Ca^2+^ stimulate TseH-mediated killing against *E. coli*

To determine the difference between agars that may account for the TseH killing efficiency, we first used XRF (X-Ray Fluorescence) to compare element contents in these two agar samples. XRF results show that Ca and Mg content were substantially higher in source 2 agar (Supplementary Table 1). Using ICP-MS (Inductively Coupled Plasma Mass Spectrometry), we further quantified Mg and Ca content. The Mg and Ca content were 85.83±2.40 mg/kg and 16.92±0.30 mg/kg respectively in source 1 agar. However, in source 2 agar, the Mg and Ca content were 515.29±13.57 mg/kg and 200.93±9.29 mg/kg (Table 1), about 6-fold and 10-fold higher than their levels in source 1 agar, respectively.

**Table 1.** Determination of Mg and Ca in agar powder by ICP-MS

|  | Mg (mg/kg) | Ca (mg/kg) |
| --- | --- | --- |
| Agar (Source 1) | 85.83±2.40 | 16.92±0.30 |
| Agar (Source 2) | 515.29±13.57 | 200.93±9.29 |

To test if the observed high levels of Mg and Ca are responsible for the enhanced TseH-mediated killing against *E. coli*, we supplemented Mg^2+^ or Ca^2+^ in source 1 agar to the detected levels and tested their effects on bacterial competition. Indeed, the addition of Mg^2+^ or Ca^2+^ stimulated TseH-mediated killing and reduced *E. coli* survival significantly, with Ca^2+^ showing a stronger effect (Figure 1B, Supplementary Figure 1B). Notably, the combination of 0.27 mM Mg^2+^ and 0.07 mM Ca^2+^, the amount closely resembling the detected levels in source 2 agar (Table 2), did not fully reduce *E. coli* survival to that in source 2 agar, suggesting other factors in source 2 also contribute to the increased TseH toxicity.

**Table 2.**
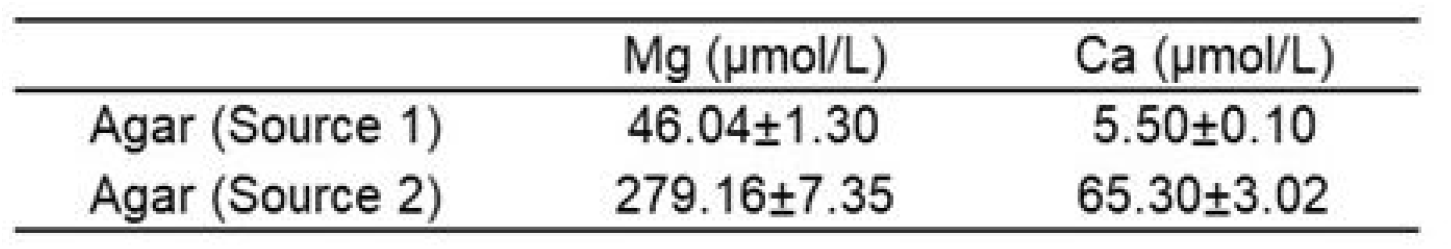
Content of Mg and Ca in LB plates with 1.3% agar

To further confirm that metal ions play a critical role in stimulating TseH-mediated killing against *E. coli*, we washed the source 2 agar powder with a metal-chelator EDTA solution to remove divalent cations. Results showed that EDTA-washed source 2 agar failed to support TseH-mediated killing, while the deionized water (ddH_2_O)-treated source 2 agar could still stimulate TseH killing (Figure 1C, Supplementary Figure 1C). Both the EDTA-treatment and water-treatment did not support TseH-mediated killing on source 1 agar plates.

Finally, we also used the defined M9 medium for competition between the *tseH*^*+*^ mutant and *E. coli*. Both M9 with source 1 agar and source 2 agar plates supported TseH-mediated killing, with the source 2 agar plate showing a stronger effect (Figure 1D, Supplementary Figure 1D). Because the M9 salt contains Mg^2+^ and Ca^2+^(1.0 mM Ca^2+^, 0.1 mM Mg^2+^), we also tested the effects when these two cations were omitted. Results showed that only source 2 agar supported the TseH-mediated killing. Collectively, these results demonstrated that TseH-mediated toxicity is largely dependent on environmental Mg^2+^ and Ca^2+^ levels.

### Conditional toxicity of TseH is also appliable to *V. cholerae*

We have previously reported that deletion of *tseH* cognate immunity gene, *tsiH*, did not render the *V. cholerae* mutant susceptible to TseH [40], which is attributed to protection by immunity-independent defense mechanisms, such as the WigKR two-component system [18]. Since *E. coli* showed different sensitivity against TseH with or without Mg^2+^ and Ca^2+^, we asked whether the *V. cholerae ΔtsiH* mutant would be sensitive to TseH in the presence of these two cations. We constructed a *ΔtsiH* deletion mutant lacking the *paar2*-*tseH*-*tsiH* three-gene operon in the *4eff*_*c*_ mutant background and used it as prey for competition against the *tseH*^*+*^ strain or the *4eff*_*c*_ strain, separately. The competition assay results showed that, like *E. coli*, the *ΔtsiH* mutant was sensitive to the *tseH*^*+*^ strain on source 2 agar plates but not on source 1 agar plates, exhibiting a 10-fold reduced survival relative to that exposed to the *4eff*_*c*_ killer strain (Figure 1E, Supplementary Figure 1E). In addition, supplementation of 0.3 mM Ca^2+^ to source 1 agar reduced the *ΔtsiH* mutant survival significantly, while a combination of 0.27 mM Mg^2+^ and 0.07 mM Ca^2+^ or 0.3 mM Mg^2+^ alone had minor effects. Thus, Ca^2+^ also stimulated TseH-mediated killing against the *V. cholerae ΔtsiH* mutant.

### The addition of Mg^2+^ and Ca^2+^does not affect T6SS secretion

Intrigued by the stimulating effect of Mg^2+^ and Ca^2+^ on TseH-mediated killing, we first hypothesized that Mg^2+^ and Ca^2+^ may affect T6SS secretion efficiency. Thus, we compared the secretion of Hcp with or without Mg^2+^ and Ca^2+^ as an indicator for T6SS activities. The results showed that Hcp was secreted to the same levels with or without Mg^2+^ and Ca^2+^ supplementation between wild type or the *tseH*^+^ samples, suggesting the addition of Mg^2+^ and Ca^2+^ has little effect on the T6SS secretion (Figure 2A, Supplementary Figure2A).

**Figure 2.**
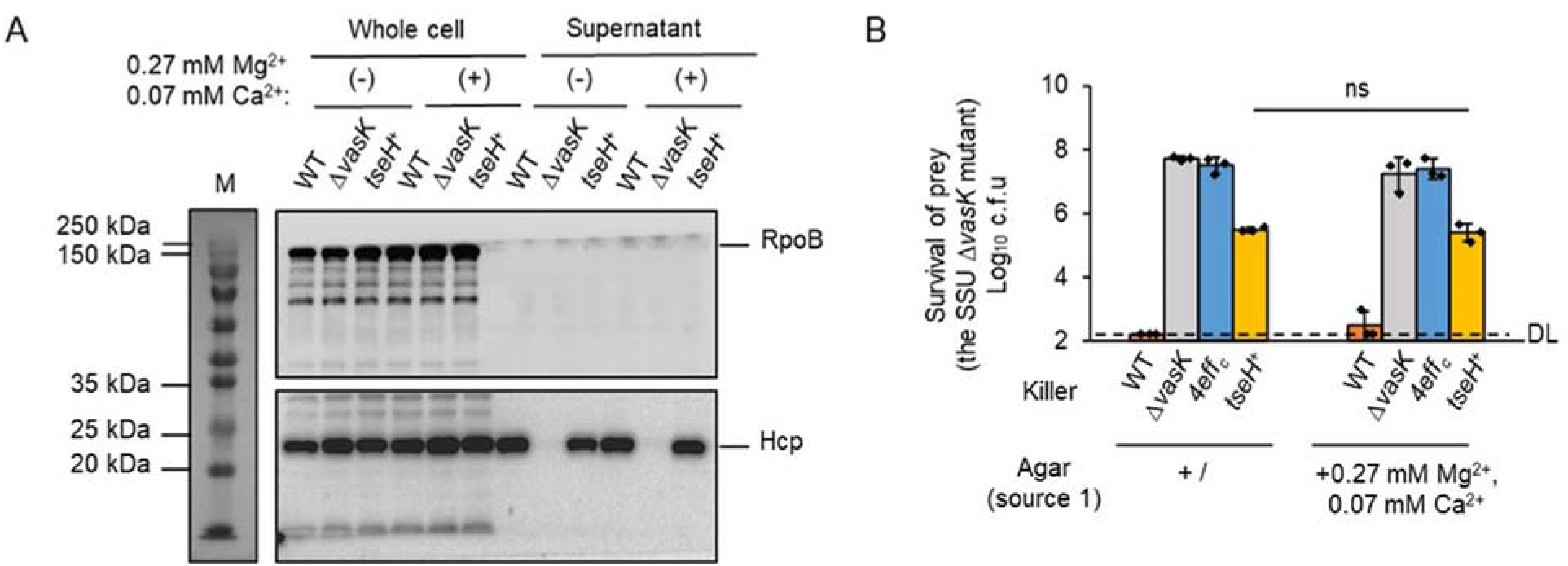
Addition of Mg^2+^ and Ca^2+^ does not affect T6SS secretion. **A**, Secretion analysis of Hcp in *V. cholerae* strains with or without Mg^2+^ and Ca^2+^. WT, wild type; Δ*vasK*, the T6SS-null Δ*vasK* mutant; *tseH*^*+*^, the *tseH*^*+*^ only mutant. (-), no cations addition; (+), addition of 0.27 mM Mg^2+^ and 0.07 mM Ca^2+^. RpoB, the beta subunit of DNA-directed RNA polymerase, serves as cell autolysis and loading control. Full images were shown in **Supplementary Figure 2A. B**, Survival of the *Aeromonas dhakensis* SSU *ΔvasK* mutant after competition with *V. cholerae* strains, as indicated. WT, wild type; Δ*vasK*, the T6SS-null Δ*vasK* mutant; *4eff*_*c*,_ the 4-antibacterial-effector-inactive mutant; *tseH*^*+*^, the TseH-active mutant. Survival of prey strains was enumerated by serial plating on selective medium. Killer survival was shown in **Supplementary Figure 2B**. Error bars indicate the mean ± standard deviation of three biological replicates. Statistical significance was calculated using a two-tailed Student’s *t*-test. ns, not significant; **, *p* <0.01. DL, detection limit.

Next we tested whether TseH requires Mg^2+^ and Ca^2+^ for its activity. TseH is shown to be inactive *in vitro*, and the structural analysis reveals that a short loop might block substrate entry [18]. Therefore, we used an *in vivo* assay by testing TseH toxicity in sensitive prey *A. dhakensis* [18]. Results showed that, with and without Mg^2+^ and Ca^2+^, the survival of *A. dhakensis* was equally reduced about 2-logs by the *tseH*^+^ strain (Figure 2B, Supplementary Figure 2B), relative to that competed with the *4eff*_*c*_ and Δ*vasK* mutant, suggesting that TseH toxicity is independent of Mg^2+^ and Ca^2+^. Taken together, these results indicated that the metal-dependent sensitivity of *E. coli* to TseH was not caused by difference in *V. cholerae* T6SS secretion nor changed TseH activities under these two conditions.

### PhoPQ two-component system contributes to immunity-independent defense against TseH

Next, we tested whether sensitivity to TseH results from metal-related defense response in *E. coli*. The PhoPQ two-component system is a key regulatory system and activated in response to Mg^2+^ starvation in *E. coli* [42, 43]. Since the TseH-mediated killing was only observed at high Mg^2+^ / Ca^2+^ levels, we speculated that the PhoPQ system is involved in modulating TseH-killing effects. Specifically, under low Mg^2+^ conditions, the PhoPQ may be activated to confer protection against TseH, while the supplement of Mg^2+^ represses such protection.

To test this hypothesis, we constructed the *E. coli* Δ*phoQ* deletion mutant by homologous recombination and used competition assay to compare its survival against the *tseH*^*+*^ strain. Results showed that the deletion of *phoQ* made *E. coli* significantly more sensitive to TseH even in the absence of Mg^2+^ and Ca^2+^ (Figure 3A, Supplementary Figure 3A). We observed a less than one-log survival difference between *E. coli* wild-type and the Δ*phoQ* strains when competed with the *tseH*^*+*^ mutant with Mg^2+^ and Ca^2+^. We also constructed the *E. coli phoQ*^*D179L*^ strain, a PhoQ locked-on mutant that constitutively phosphorylates PhoP and activates its regulon genes [44]. Competition analysis showed that this *phoQ*^*D179L*^ mutation significantly increased *E. coli* survival relative to the Δ*phoQ* with or without Mg^2+^ and Ca^2+^ (Figure 3A). These results suggest that the PhoPQ two-component system also contribute to immunity-independent defense mechanism against TseH toxicities, in addition to the known envelope stress response systems [18].

**Figure 3.**
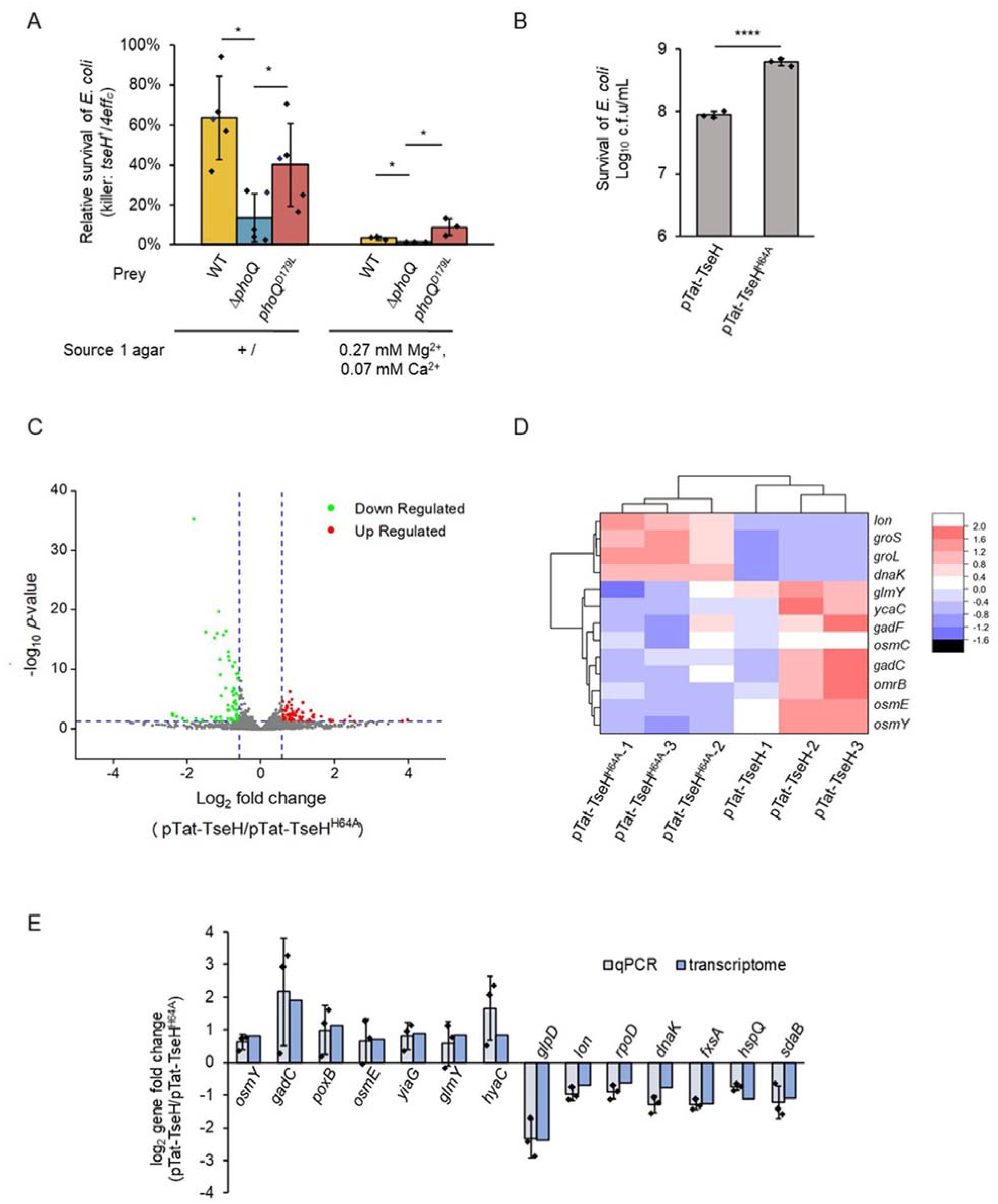
Innate immunity-like pathways protect *E. coli* against TseH. **A**, Relative survival of *E. coli* strains competed with the *V. cholerae 4eff*_*c*_ and *tseH*^*+*^ strain (*tseH*^*+*^/*4eff*_*c*_). WT, wild-type; Δ*phoQ*, the *phoQ* deletion mutant; *phoQ*^*D179L*^, a constitutively active *phoQ* mutant. Survival of prey strains were enumerated by serial plating on selective medium. Killer survival was shown in **Supplementary Figure 3A. B**, Survival of *E. coli* ectopically expressing TseH and its inactive mutant TseH^H64A^ with a periplasmic localization Tat signal. Cells were induced with 0.1% [w/v] arabinose and survival was enumerated by serial plating on plates with 0.2% [w/v] glucose. **C**, Volcano plot of RNA-seq results in *E. coli* samples ectopically expressing Tat-TseH and its inactive mutant Tat-TseH^H64A^. Differential expression genes were screened by setting cut-off value with fold change > 1.5, *p*-value < 0.05 and an average FPKM >10. Up and down-regulated genes were indicated in red and green, respectively. Genes with no significant change were indicated in gray. **D**, Cluster analysis and heat map of differential expression mRNAs in *E. coli* samples ectopically expressing Tat-TseH and its inactive mutant Tat-TseH^H64A^. Scale means the normalized FPKM of each sample by Z-score. **E**, qRT-PCR verification of differential expression of mRNA genes. The 16S rRNA gene serves as the reference. For **A, B** and **E**, error bars indicate the mean ± standard deviation of at least three biological replicates. Statistical significance was calculated using a two-tailed Student’s *t*-test. *, *p* <0.05; ****, *p*< 0.0001. DL, detection limit.

### Transcriptome analysis in *E. coli* periplasmically expressing TseH and its inactive mutant TseH^H64A^

Next, we examined the effect of adding metal cations to *E. coli* using RNA-seq transcriptome analysis. By transferring and incubating exponential-phase-growing *E. coli* to LB source-1-agar plates with and without 0.3 mM Mg^2+^ for 30 min prior to RNA extraction, a condition mimicking competition assays, we found very few significant changes for the transcriptomes under these two conditions (Supplementary Figure 3B, Dataset 1). Then, we examined the effect of TseH toxicity on transcriptome expression by ectopically expressing Tat-TseH and its inactive mutant Tat-TseH^H64A^ using arabinose-inducible pBAD vectors in *E. coli*. First, we established the induction condition to be 30-min so that the survival of *E. coli* was reduced by TseH to 14.5% of the *E. coli* survival expressing the non-toxic mutant (Figure 3B), suggesting TseH was expressed in most cells. Under this condition, we performed RNA extraction and performed transcriptome analysis. To identify genes differentially expressed, we set the cut-off value of fold change > 1.5 and *p*-value < 0.05. We also removed the low transcriptome gene with an average FPKM (fragments per kilobase per million mapped fragments) < 10. By comparing the transcriptome of these two strains, we found 106 genes were differentially expressed, of which 50 genes were up-regulated and 56 genes were down-regulated (Figure 3C, Supplementary Figure 3C, Dataset 2).

Among the up-regulated genes, 22 genes belonged to the RpoS general stress response regulon, including *gadCF* and *osmY* (Figure 3D). We also found three small regulatory RNA, *gadF, glmY* and *omrB* were up-regulated. *GadF* is involved in acid stress response [45, 46]; *GlmY* controls the amino sugar metabolism in *E. coli* by post-transcription and deletion of *glmY* renders cells sensitive to envelope stress [47]. *OmrB* is a small RNA that is involved in regulating the protein composition of the outer membrane [48]. Among the down-regulated genes, we found 24 genes belonging to the heat-shock response RpoH regulon were down-regulated, including *dnaK, groL, groS*, and the protease gene *lon*. These genes are important for proper protein folding.

We have shown that the T6SS delivered-TseH induces the envelope stress response in *E. coli* [18]. In the transcriptome results, we found the BaeR regulon gene *ycaC* was up-regulated. In addition, *gadF*, also a PhoPQ-regulated gene, was upregulated. Another PhoPQ-regulated gene *mgrB*, involved in Mg^2+^ uptake, was up-regulated with *p*-value<0.05 but with a moderate fold change <1.5.

To validate the RNA-seq data, we performed qRT-PCR assays of 14 genes, of which 7 were up-regulated and 7 were down-regulated. These genes were with the relative high fold change in RNA-seq results. The 16S rRNA gene was used as the reference. The qRT-PCR results were consistent with the RNA-seq results (Figure 3E).

### Divalent cations contribute to TseL-mediated toxicity

We also noticed that another T6SS effector TseL with phospholipase activity in *V. cholerae*, also exhibited different toxicity against *E. coli* on different agar sources during interspecies competition. On source 1 agar plates, the *V. cholerae tseL*^*+*^ mutant reduced *E. coli* survival by 100-fold relative to the *4eff*_*c*_ mutant. In contrast, on source 2 agar plates, the *E. coli* survival was reduced about 10,000-fold by the *tseL*^*+*^ mutant relative to the *4eff*_*c*_ mutant (Figure 4A, Supplementary Figure 4A). In addition, EDTA-treatment of different agar sources also abolished TseL-mediated killing difference between the two agar sources (Figure 4B, Supplementary Figure 4B). When comparing *E. coli* survival on TseL-sensitive agar source 2, we found that EDTA-treatment increased *E. coli* survival by 100-fold relative to deionized water-treatment. To further investigate which divalent cations play a role in this process, we competed the *tseL*^*+*^ mutant and *E. coli* on source 1 agar plates supplemented with 0.3 mM Mg^2+^, Ca^2+^, Ni^2+^ and Cu^2+^, respectively. However, unlike TseH, none of the tested cations increased the TseL-toxicity to the agar source 2 levels (Figure 4C, Supplementary Figure 4C). Nonetheless, these results suggest that TseL-mediated toxicity is also modulated by some EDTA-chelatable divalent cations.

**Figure 4.**
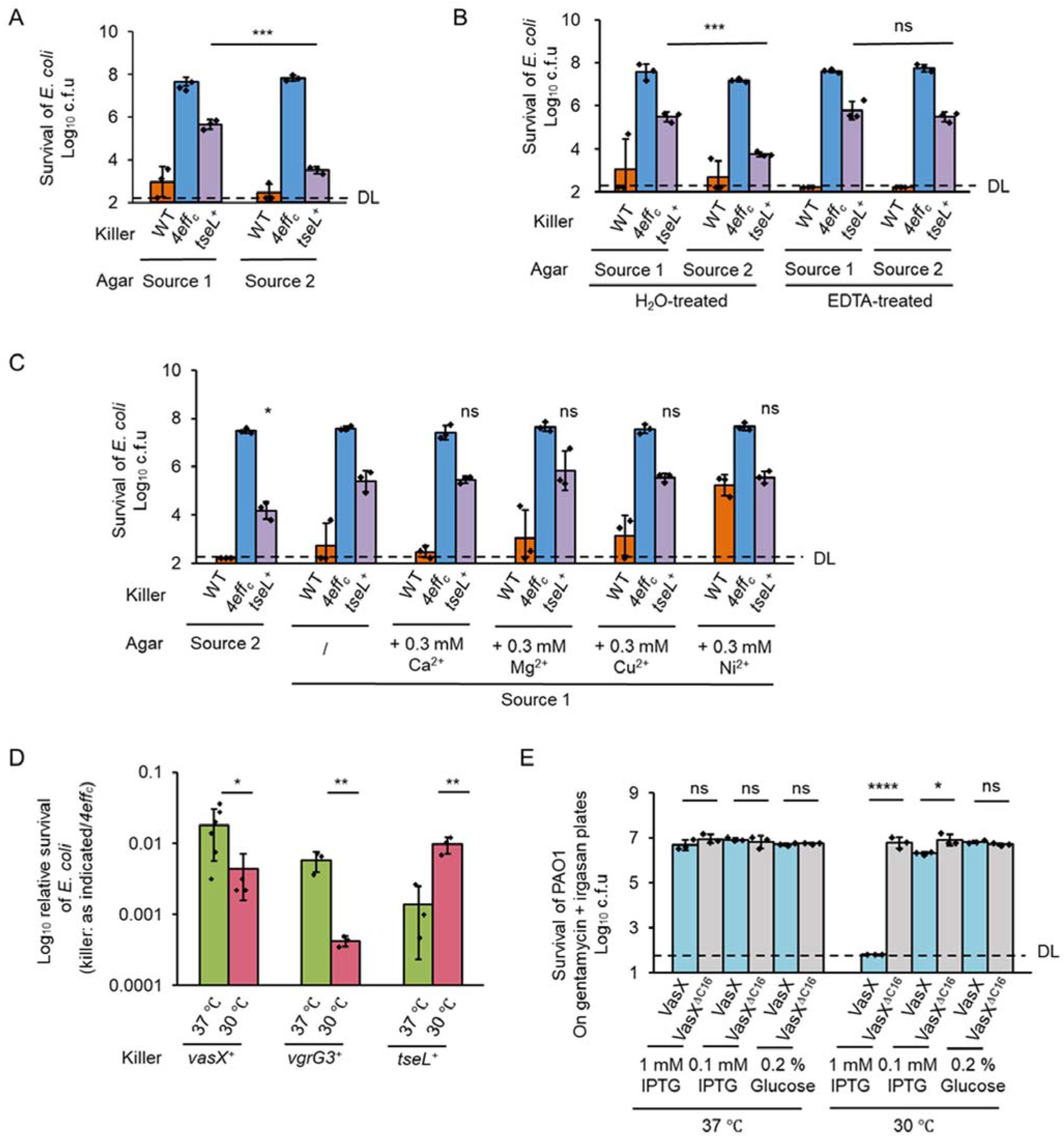
Effects of cations and temperature on killing by other effectors. **A**, Survival of *E. coli* after competition with *V. cholerae* TseL-active mutant *tseL*^*+*^ on LB source-1-agar or source-2-agar plates. **B**, Effect of EDTA-treatment on survival of *E. coli* competed with *V. cholerae tseL*^*+*^. The ddH_2_O-treated agar plates serve as control. **C**, Effect of cation supplementation on TseL-mediated *E. coli* killing. LB source-1-agar plates were supplemented with 0.3 mM Ca^2+^, Mg^2+^, Cu^2+^ or Ni^2+^, respectively. **D**, Relative survival of *E. coli* after competed with *V. cholerae* strains at 30 °C or 37 °C (normalized to *4eff*_*c*_), as indicated. For **A** to **D**, WT, wild type; *4eff*_*c*_, the 4-antibacterial-effector-inactive mutant; *tseL*^*+*^, the TseL-active only mutant. *vasX*^*+*^, the VasX-active only mutant. *vgrG3*^*+*^, the VgrG3-active only mutant. Survival of *E. coli* strains was enumerated by serial plating on selective medium. Survival of killer strains was shown in **Supplementary Figure 4A-4D**, respectively. **E**, Toxicity assay of *P. aeruginosa* PAO1 strains ectopically expressing Tat-VasX and its colicin-inactivated mutant Tat-VasX^ΔC16^ on plates with gentamycin and irgasan at 30 °C or 37 °C. For **A-E**, error bars indicate the mean ± standard deviation of three biological replicates. Statistical significance was calculated using a two-tailed Student’s *t*-test for two groups comparison or one-way ANOVA test for more than two groups comparison. *, *p* <0.05; **, *p* <0.01; ****, *p*< 0.0001. DL, detection limit.

### Temperature affects effector-mediated competition

Since *V. cholerae* may also experience temperature changes in the environment or during transmission from environment to the host, we next determined whether temperature also plays a role in effector-mediated bacterial competition. We used a panel of *V. cholerae* single-effector-active killer strains and *E. coli* as prey. Results show that the relative survival of *E. coli* was significantly reduced at 30 °C in comparison to that at 37 °C when *E. coli* was competed with the *vasX*^*+*^ (with membrane pore-forming activity) and the *vgrG3*^*+*^ (a lysozyme) strains, separately (Figure 4D, Supplementary Figure 4D). In contrast, the survival of *E. coli* was significantly increased at 30 °C relative to that at 37 °C when *E. coli* was competed with the *tseL*^*+*^ strain (Figure 4D, Supplementary Figure 4D). These results collectively indicate that temperature also plays an important role in T6SS-mediated competition.

### VasX-toxicity in *P. aeruginosa* is dependent on temperature and the antibiotic irgasan

We have previously reported that periplasmic expression of VasX using a Tat secretion signal is highly toxic in *E. coli* but not in *Pseudomonas aeruginosa* PAO1, an important opportunistic pathogen [19]. We thus tested whether the resistance of *P. aeruginosa* to VasX is also temperature-dependent. First, we confirmed this resistance phenotype by comparing the survival of PAO1 expressing plasmid-borne Tat-VasX or its inactive mutant Tat-VasX^ΔC16^ [17] at 37 °C (Figure 4E). However, when VasX was highly induced at 30 °C, we found that survival of PAO1 was reduced to undetectable levels (Figure 4E). Because the assays were performed on plates containing two antibiotics, irgasan that PAO1 is intrinsically resistant to, and gentamycin whose resistance is conferred by the pTat-VasX plasmid, and because expression of VasX increases membrane permeability [19], we hypothesized that the increased sensitivity to VasX may be caused by the presence of antibiotics. We repeated periplasmic expression of VasX in PAO1 but only plated the induced cells on gentamycin. We found that PAO1 became resistant to VasX expression at both 30 °C and 37 °C (Supplementary Figure 4E). These results suggest that the membrane damages caused by VasX expression at 30 °C, albeit insufficient to cause cell death directly, can disturb the intrinsic resistance of *P. aeruginosa* to irgasan (Figure 4E, Supplementary Figure 4E).

## Discussion

Microbes have been found in almost all ecological niches on earth and contribute to a variety of functions ranging from macroscale carbon recycling and waste removal to individual-level host health and disease. These functions are often determined by not a single but multiple species that coexist in complex communities, with extensive interspecies interactions via various molecular mechanisms. The T6SS is one such crucial mechanism exhibiting anti-bacterial, anti-fungal and other anti-eukaryotic functions [5, 24, 49–51]. From a molecular ecological perspective, the T6SS-mediated interspecies competition is a complex and multifaced process whose outcome is determined by diverse factors including the number and type of the secreted effectors [31], the frequency of T6SS firing correlated with energy state [31], the immunity protein-dependent specific protection [24], the non-specific stress response pathways in killer and prey cells [18, 20], and the availability of nutrients [52][53]. Here, we add on to this list by showing that abiotic environmental factors can modulate prey cell sensitivity to effector toxicities, thereby affecting T6SS-mediated competition (Figure 6). Our results demonstrate not only that some seemingly inactive effectors may be highly toxic in natural settings but also the importance of examining toxin-mediated interspecies interactions beyond the routine lab conditions.

Almost all T6SS species secrete multiple effectors with obvious benefits for T6SS-mediated competitive fitness [31, 37, 39]. However, many effectors are phenotypically inactive, hindering the full understanding of the ecological functions of the T6SS. An elegant strategy employing barcoding-sequencing analysis of mixed effector-susceptible populations in *P. aeruginosa* reveals that effectors may act jointly with other effectors to exert synergistic functions but their activities can also be variable depending on salinity, pH and aerobic conditions [39]. Here we show the ecologically-relevant abiotic factors, including cations and temperature, that microbes frequently encounter in nature environment, play a critical role in mediating T6SS-dependent interspecies killing. Our previous study on the *V. cholerae* T6SS effector TseH reveals that *E. coli* mutants impaired in envelope stress response (ESR) pathways, but not the wild type, are sensitive to TseH toxicities [18]. Here we show that supplementation of Mg^2+^ and Ca^2+^ cations can render *E. coli* susceptible to TseH despite of the intact ESR protection. Another effector TseL has also exhibited cation-related toxicity since EDTA-treatment of the agar could reduce TseL-mediated killing against *E. coli*. Notably, as *V. cholerae* is a waterborne pathogen commonly found in water sources in which Mg^2+^ (average concentration 50 mM) and Ca^2+^ (∼ 10 mM) are present at much higher levels than the levels supplemented here [54], the *V. cholerae* T6SS effectors may have a broader target range in the natural niche. Indeed, when a source of raw agar was used, we also found highly increased TseH-mediated killing against *E. coli* (Figure 5A, 5B, Supplementary Figure 5A, 5B).

**Figure 5.**
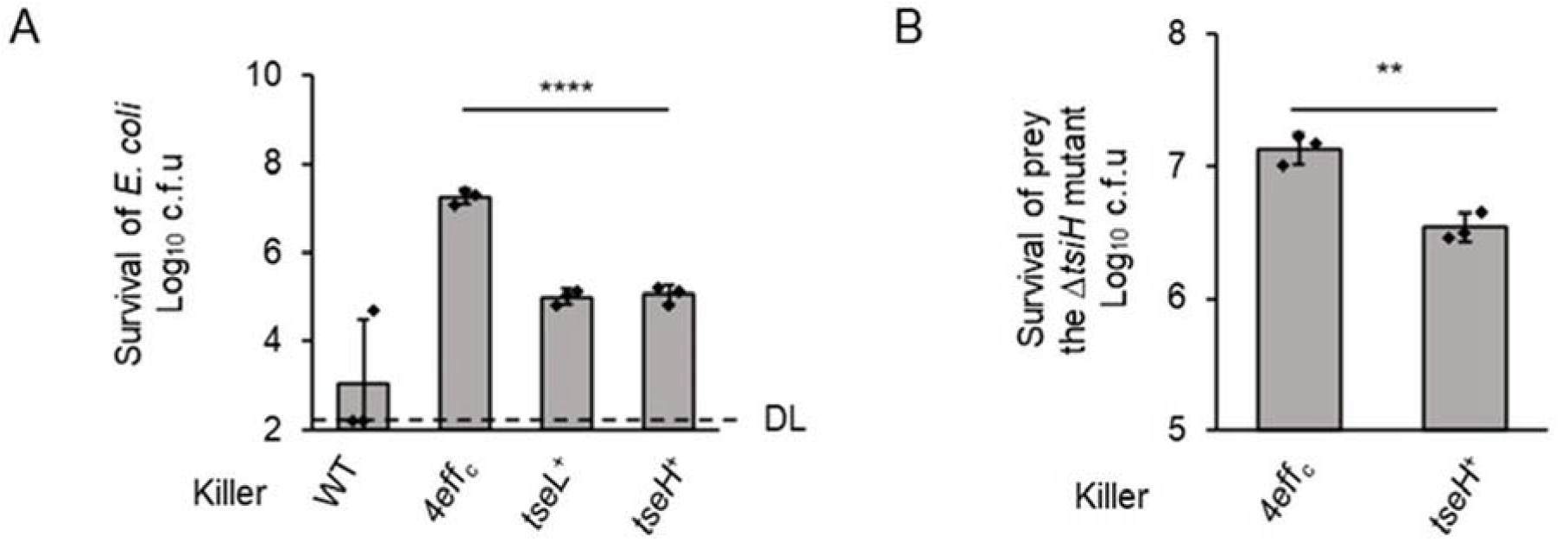
Competition analysis of TseH-mediated killing on raw agar. **A**, Survival of *E. coli* prey competed with *V. cholerae* strains on LB raw agar plates, as indicated. **B**, Survival of the *V. cholerae ΔtsiH* mutant as prey. For **A** and **B**, killer strains are indicated at the bottom. WT, wild type; *4eff*_*c*,_ the 4-antibacterial-effector-inactive mutant; *tseL*^*+*^, the TseL-active only mutant. Survival of prey cells was enumerated by serial plating on selective medium. Killer survival was shown in **Supplementary Figure 5A-5B** respectively. Error bars indicate the mean ± standard deviation of three biological replicates. Statistical significance was calculated using a two -tailed Student’s *t*-test. **, *p* <0.01; ****, *p*< 0.0001. DL, detection limit.

**Figure 6.**
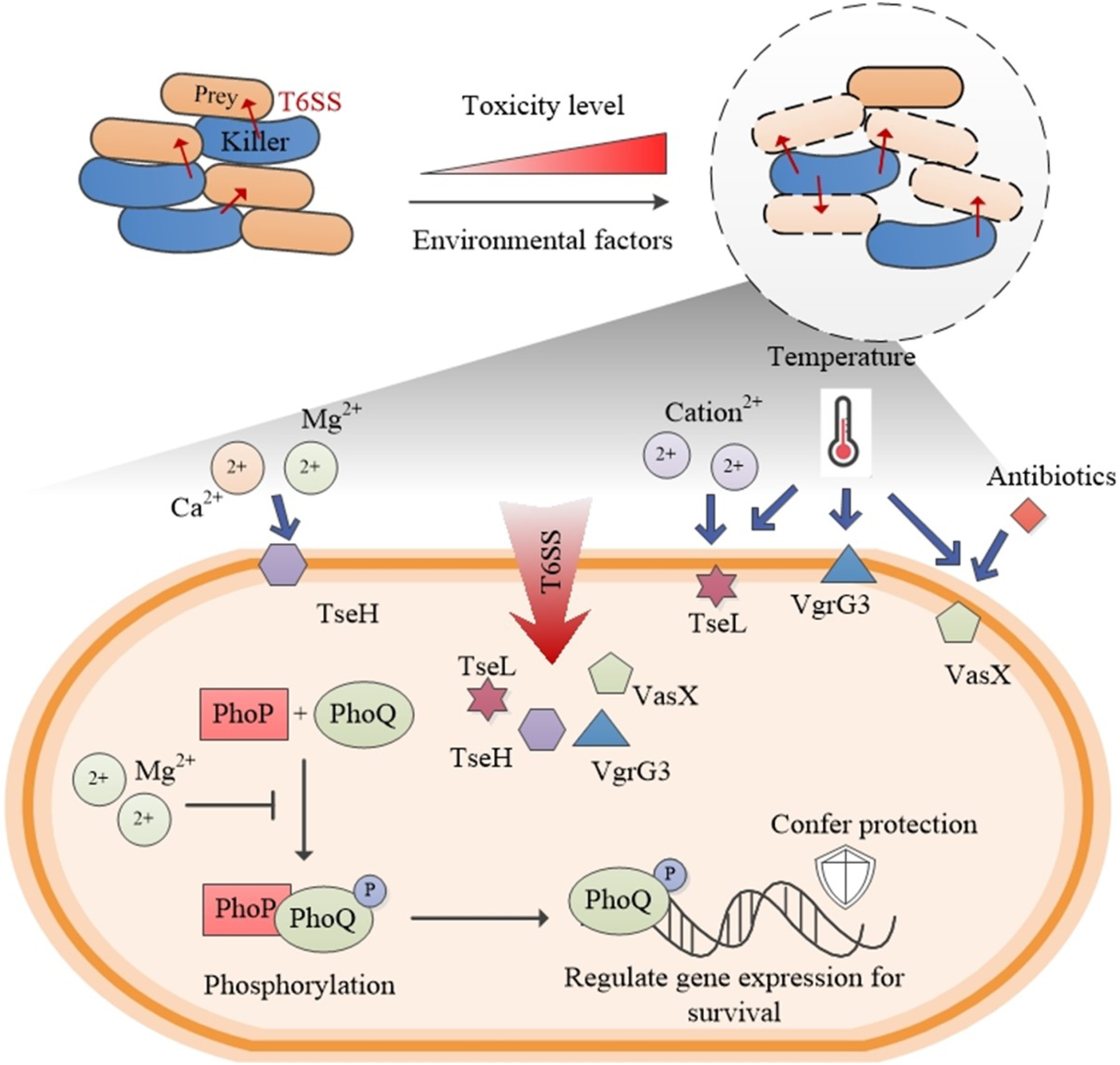
Abiotic factors and stress response dictate the outcome of T6SS-mediated competition. Environmental abiotic factors affect the susceptibility of prey cells to T6SS effectors. These factors include the presence metal cations, temperature, antibiotic molecules. Prey cells that do not possess specific immunity proteins rely on innate immunity-like stress response pathways for protection. As an example, the PhoPQ two-component system was activated under Mg^2+^-limited environment, which is important for protecting *E. coli* from TseH-mediated toxicity. Temperature and other environmental factors likely also modulate prey cell physiology to alter cell sensitivity to T6SS-mediated competition. These abiotic factors thus play a key role in the dynamic competition of a T6SS-related microbial community.

Mg^2+^ and Ca^2+^ are known to have critical cellular functions, but the effect of cations on T6SS effector toxicities in prey cells has not been reported before. As the most abundant divalent cation in living cells, Mg^2+^ neutralizes the negative charge of biomolecules, including nucleotides and nucleic acids, phospholipids in various membrane structures in cells, as well as serving as an indispensable cofactor in various enzymes [41, 55, 56]. Like Mg^2+^, Ca^2+^ could also maintain cell structure, participate in motility and cell division processes [57, 58]. Due to the critical role Mg^2+^ and Ca^2+^ play in bacteria, the concentrations of Mg^2+^ and Ca^2+^ are tightly regulated [56, 59]. Under low Mg^2+^ conditions, the two component system PhoPQ is activated to regulate a large number of genes involved in virulence, metal uptake, and other important functions [41, 56]. The increased resistance of *E. coli* to TseH in agar source 1 with lower Mg^2+^ may thus be attributed to activation of PhoPQ. Indeed, deletion of *phoQ* leads to increased sensitivity to TseH, suggesting that the PhoPQ-system is important for such protection (Figure 3A).

Although we have yet to determine the exact physiological changes in prey cells that account for the temperature and cation-dependent effector-mediated competition, it is likely a multifaceted process that involves a number of different cellular pathways, as exemplified in the transcriptome changes upon TseH induction (Figure 3C, 3D). Given the prevalence of T6SS organisms, including plant and host pathogens, as well as the diversity of effectors in each species, it is crucial to take into account of ecologically relevant conditions for testing T6SS-related effector functions and interspecies interactions in future studies. The environmental cues, such as cations and temperature, may have profound effects on the composition of T6SS-containing microbial communities and the evolution of their defense and attack mechanisms.

## Methods

### Strains and growth conditions

Strains and plasmids used in this study are described in Supplementary Table 2. All constructs were verified by sequencing. Primers are available in Dataset 3. Bacteria were grown in LB([w/v] 1% tryptone, 0.5% yeast extract, 0.5% NaCl) at 37 °C aerobically. Antibiotics and inducers were used at following concentrations: kanamycin (25 µg/mL for *E. coli*, 50 µg/m L for *V. cholerae* and *A. dhakensis*), streptomycin (100 µg/mL), chloramphenicol (2.5 µg/mL for *V. cholerae* and 25 µg/mL for *E. coli*), irgasan (25 µg/mL), gentamycin (20 µg/mL), L-arabinose ([w/v] 0.1% or 0.01% as indicated), IPTG (0.1 mM or 1 mM as indicated). Tryptone (BIO BASIC, TG217(G211)), yeast extract (BIO BASIC, G0961), source 1 agar (BIO BASIC, FB0010), source 2 Agar (Sangon Biotech, A505255-0250).

### EDTA treatment of agar powder

Agar powders were washed by 0.1 M EDTA solution for 5 min with vortex, followed by deionized water washing with 5 times to remove EDTA. Then the treated agar was dried and could be used for bacterial competition experiments.

### Bacterial competition assay

Competing strains were grown overnight in LB with appropriate antibiotics. Killer cells were then sub-cultured to OD_600_=1. For interspecies competition, killer and prey cells were mixed at a ratio of 5:1 and co-incubated on LB-agar plates or M9-agar (93.0 mM Na^+^, 22.1 mM K^+^, 18.7 mM NH_4_^+^, 1.0 mM Ca^2+^, 0.1 mM Mg^2+^, 29.2 mM Cl^-^, 0.1 mM SO_4_^2−^, 42.2 mM PO_4_^2−^, [w/v] 0.1% glucose) plates at 37 °C or 30 °C (as indicated) for 3 h. For intraspecies competition, killer and prey cells were mixed at a ratio of 20:1 and co-incubated on LB-agar plates at 37 °C for 20 h (or 3 h when using raw agar). After co-incubation, the survival of killer and prey cells were enumerated by 10-fold serial plating on LB plates with selective antibiotics. The mean Log_10_ c.f.u of the recovered killer and prey strains was plotted and error bars show the mean ± standard deviation of at least three biological replicates. A two-tailed Student’s *t*-test and one-way ANOVA test was used to determine *p*-values.

### XRF and ICP analysis

XRF (X-Ray Fluorescence) and ICP (Inductively Coupled Plasma Mass Spectrometry) analysis was done by the Instrumental Analysis Center in Shanghai Jiao Tong university. For XRF analysis, the agar powder was pressed into disks (30 mm diameter) and performed with a XRF (XRF-1800, SHIMADZU). For ICP analysis, the two agar samples were digested with 5 mL concentrated nitric acid and diluted to 50 mL with ultrapure water. After samples were prepared, elemental analysis was performed with an ICP (Avio 500, PerkinElmer).

### Protein secretion assay

Strains were grown overnight in LB with appropriate antibiotics. Cells were then sub-cultured to OD_600_=1 with or without Mg^2+^ and Ca^2+^. 2 ml of OD_600_ 1 cells were collected by centrifuge at 2,500 × *g* for 3 min and resuspended in 1 mL fresh LB. Resuspended cells were placed at 30 °C for 1 h and centrifuged at 10,000 × *g* for 2 min at room temperature (RT). Cell pellet was used as the whole-cell sample. The supernatant was centrifuged at 10,000 × *g* for 2 min again as the secretion sample. All samples were mixed with SDS-loading dye, boiled at 98 °C for 10 min, followed by SDS-PAGE (sodium dodecyl sulfate polyacrylamide gel electrophoresis) analysis and Western blotting analysis.

### Western blotting analysis

After electrophoresis in a 12% SDS-PAGE gel, proteins were transferred to a PVDF membrane (Bio-Rad). Then, the protein-bound membrane was blocked with 5% [w/v] non-fat milk in TBST (50 mM Tris, 150 mM NaCl, 0.1% [v/v] Tween-20, pH 7.6) buffer for 1 h at RT. After blocking, the membrane was sequentially incubated with primary and secondary HRP (horseradish peroxidase)-conjugated antibodies in TBST with 1% [w/v] milk for 1 h at RT. Signals were detected using the Clarity ECL solution (Bio-Rad). The monoclonal antibody to RpoB, the beta subunit of RNA polymerase, were purchased from Biolegend (RpoB, Product # 663905). The polyclonal antibody to Hcp was custom-made by Shanghai Youlong Biotech. The HRP-linked secondary antibodies were purchased from ZSGB-Bio (Product # ZB-2305 (mouse) and # ZB-2301 (rabbit)). The Hcp antibody was used at 1:10000 dilution, while others at 1:20000 dilution.

### Toxicity assay

For VasX toxicity assay in PAO1, overnight cells carrying pPSV37 constructs were grown in LB with appropriate antibiotics and 0.2% [w/v] glucose at 37 °C. Cells were then collected and resuspended in fresh LB. A serial dilution was plated on LB plates containing 0.1 mM, 1 mM IPTG or 0.2% [w/v] glucose with antibiotics as indicated.

### RNA sample preparation

For transcriptome analysis of *E. coli* with or without Mg^2+^, exponential-phase-growing *E. coli* cells were collected and transferred onto LB plates with or without 0.3 mM Mg^2+^ for 30 min prior to RNA extraction. For transcriptome analysis of *E. coli* ectopically expressing Tat-TseH and its inactive mutant Tat-TseH^H64A^, *E. coli* strains with pBAD24kan-Tat-TseH/TseH^H64A^ plasmid were grown overnight on LB plates with appropriate antibiotics and 0.2% [w/v] glucose. Cells were then sub-cultured to OD_600_=1, collected by centrifugation at 10,000 × *g* for 0.5 min and washed twice with fresh LB. After being recovered at 37 °C for 10 min, cells were induced with 0.1% [w/v] arabinose for 30 min followed by total RNA extraction. The survival of *E. coli* strains after induction was enumerated by 10-fold serial plating on LB plates with selective antibiotics and 0.2% [w/v] glucose.

### RNA extraction

100 μL 8×lysis buffer (8% [w/v] SDS, 16mM EDTA) was added into 700 μL culture of each sample after induction and mixed well by vortex for 5 s. Then 800 μL prewarmed hot acidic phenol (65 °C) was added and mixed by inverting immediately. Tubes were incubated at 65 °C for 5 min with mixing briefly every 1 min. After putting on ice for 10 min, the mixture was centrifuged at 13,000 × *g* for 2 min. The top supernatant was carefully transferred to a new tube and an equal-volume of absolute ethanol was added. Then the crude RNA was purified by RNA prep Pure Cell/Bacteria Kit (TIANGEN, Product #DP430) and genomic DNA was removed by DNase I (NEB, Product B0303S) treatment at 37 °C for 30 min. After DNase I treatment, RNA samples were then purified with the RNA clean Kit (TIANGEN, Product #DP421). Purified RNA was electrophoresed on 1% [w/v] agarose gel to monitor the integrity and contaminants.

### Transcriptome analysis

RNA-seq was done by Novogene. Briefly, rRNA was removed from total RNA samples by using probes. Then obtained mRNA was fragmented by divalent cations. The first strand cDNA was synthesized by M-MuLV Reverse Transcriptase using random hexamers as primers. The RNA strand was degraded by RNase H. The second strand of cDNA was synthesized using dUTP to replace dTTP in dNTP mixture. The purified double-stranded cDNA was end-repaired, adding an A tail and connecting to the sequencing adapter. USER enzyme (NEB, USA) was used to degrade the second strand of cDNA containing U. After using AMPure XP beads to screen cDNA with 370∼420 bp and PCR amplification of these fragments, the final library was obtained with another cycle of AMPure XP beads purification. The clustering of the index-coded samples was performed on a cBot Cluster Generation System following to the manufacturer’s instructions, and the sequencing was performed using the Illumina Hiseq TM 2500 platform with pair-end 150 base reads. Raw data was filtered according to the following standards: (1) removing reads with unidentified nucleotides (N); (2) removing reads with low sequencing quality (>50% bases having Phred quality scores of ≤20); (3) removing reads with the adapter. The obtained clean data were used for downstream analysis. Bowtie2-2.2.3 was used to build an index of the reference genome and align clean reads to reference genome. The gene expression level was calculated and further normalized by HTSeq v0.6.1. FPKM, fragments per kilobase of transcript sequence per millions base pairs sequenced, was used to demonstrated gene expression level here.

### Bioinformatics Analysis

Differential expression genes were identified by setting cut-off value of fold change > 1.5, *p*-value < 0.05. and average FPKM > 10. PCA analysis was performed by R package FactoMineR. Person correlation coefficient was calculated by R package WGCNA. Heat map was performed in Origin App: Heat map with Dendrogram.

### qRT-PCR Analysis

14 differential expression genes were chosen from transcriptome results. Primers for qPCR were designed by PerlPrimer. The 16S rRNA gene was used as the reference. Reverse transcription was done using PrimeScript^™^ RT reagent Kit with gDNA Eraser (Perfect Real Time) (Takara, #RR047A). qPCR reaction was prepared by TB Green® Premix Ex Taq^™^ (Tli RNaseH Plus) (Takara, #RR420A) and detected by CFX Connect Real-Time PCR Detection System (BIO-RAD, #1855201). Analysis of relative gene expression was calculated using the 2^-ΔΔCT^ method. Each sample was measured in triplicate and repeated at least three times.

## Data availability

The data that support the findings of this study are available within the paper or available from the corresponding author upon reasonable request.

## Acknowledgements

This work was supported by funding from National Key R&D Program of China (2018YFA0901200), National Natural Science Foundation of China (31770082 and 32030001), Canadian Institutes of Health Research, Natural Sciences and Engineering Research Council of Canada, and Canada Research Chair program. The funders had no role in study design, data collection and interpretation, or the decision to submit the work for publication.

## Author contributions

T.D. conceived the project. M.T. performed most of the experiment and data analysis. T.P., Z.W., and H.L. performed experiments. X.W. contributed to RNA-seq data analysis. M.T. prepared the first draft. T.D. contributed to the revision with assistance from M.T. and T.P..

**Correspondence and request for materials should be addressed to T. Dong**.

