## Supplementary Materials for "Abiotic factors modulate interspecies competition mediated by the type VI secretion system effectors in *Vibrio cholerae*"

Supplementary Figure 1 :

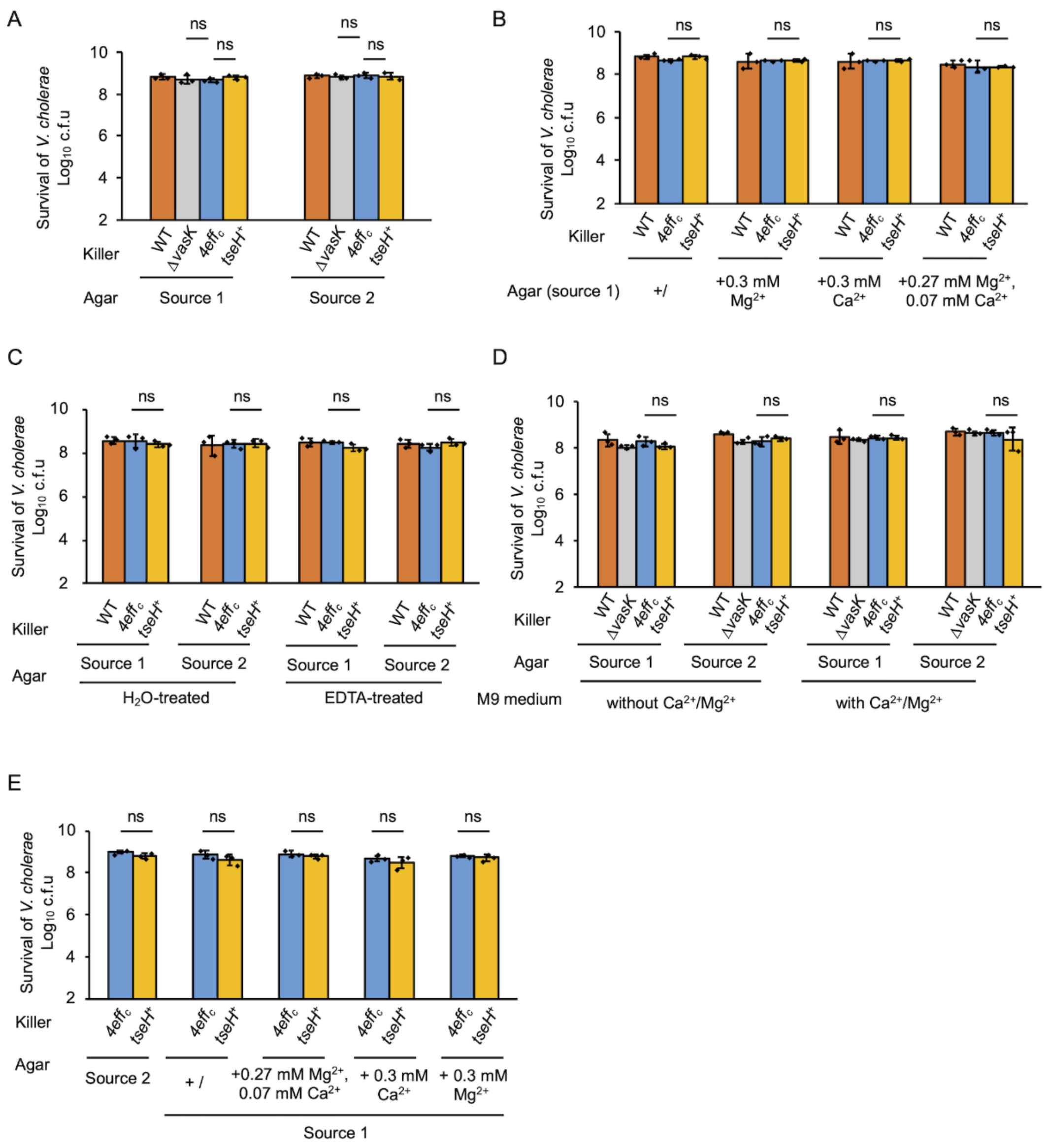


**Supplementary Figure 1. Survival of killers after competition showed equal survival, relating to Figure 1. A-D**, Survival of *V. cholerae* after competition with *E. coli* on LB source-1-agar or source-2-agar plates (**A**), on LB source-1-agar plates supplemented with $Mg^{2+}$ and $Ca^{2+}$ (**B**), on LB EDTA-treated agar plates (**C**), and on M9 source-1-agar or source-2-agar plates (**D**). **E,** Survival of *V. cholerae* after competition with the *V. cholerae* Δ*tsiH* mutant. For **A-E,** WT, wild type; Δ*vasK*, the T6SS-null Δ*vasK* mutant; $4eff_C$, the 4-antibacterial-effector-inactive mutant; $tseH^+$, the TseH-active mutant. Survival of killer strains was enumerated by serial plating on selective medium. Error bars indicate the mean ± standard deviation of three biological replicates. Statistical significance was calculated using a two-tailed Student's *t*-test for two groups comparison or one-way ANOVA test for more than two groups comparison. ns, not significant.

Supplementary Figure 2 :

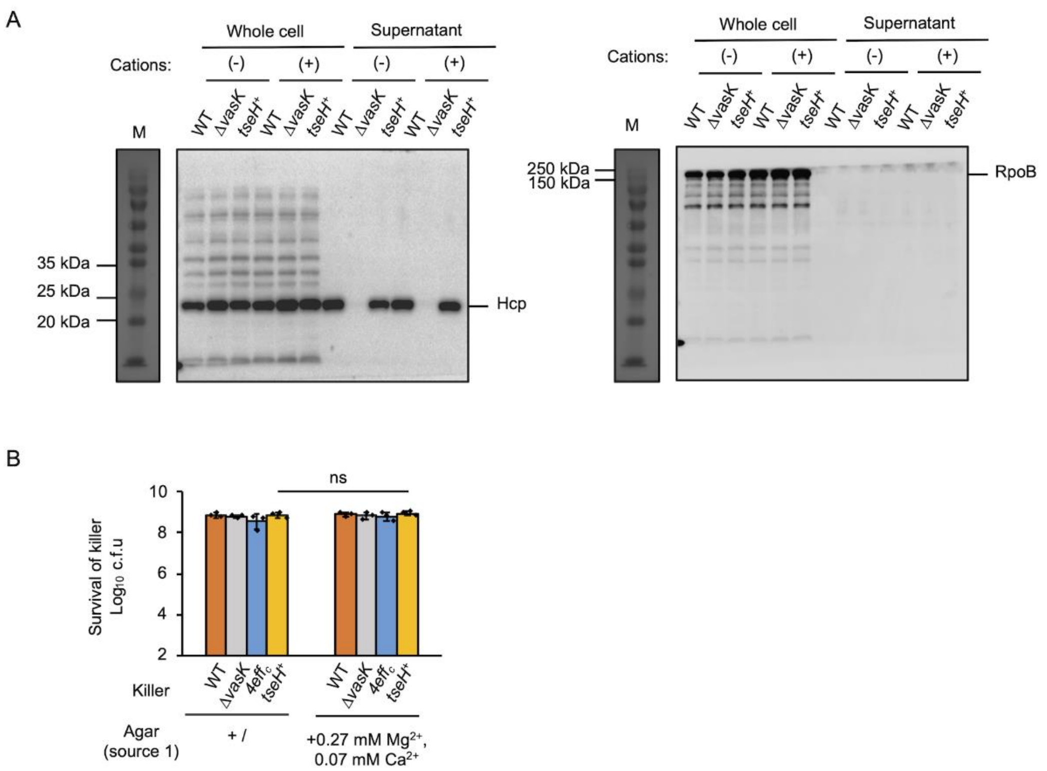


**Supplementary Figure 2. The addition of cations does not stimulate Hcp secretion or T6SS killing. A,** Full images of Hcp secretion in *V. cholerae* strains supplemented with or without $Mg^{2+}$ and $Ca^{2+}$ in **Figure 2A**. Left panel, Hcp signals. Right panel, RpoB signals. WT, wild type; Δ*vasK*, the T6SS-null Δ*vasK* mutant; $tseH^{+}$, the TseH-active only mutant. (-), no cations addition; (+) addition of 0.27 mM $Mg^{2+}$ and 0.07 mM $Ca^{2+}$. RpoB, the beta subunit of DNA-directed RNA polymerase that cannot be secreted, was used as control. **B,** Survival of killers in competition assay showed equal survival, relating to **Figure 2B**. Killer survival was selected by the streptomycin resistant on the genome. WT, wild type; Δ*vasK*, the T6SS-null Δ*vasK* mutant, $4eff_{c,}$ the 4-antibacterial-effector-inactive mutant; $tseH^{+}$, the TseH-active mutant. Error bars indicate the mean ± standard deviation of three biological replicates. Statistical significance was calculated using a two-tailed Student's *t*-test. ns, not significant.

Supplementary Figure 3：

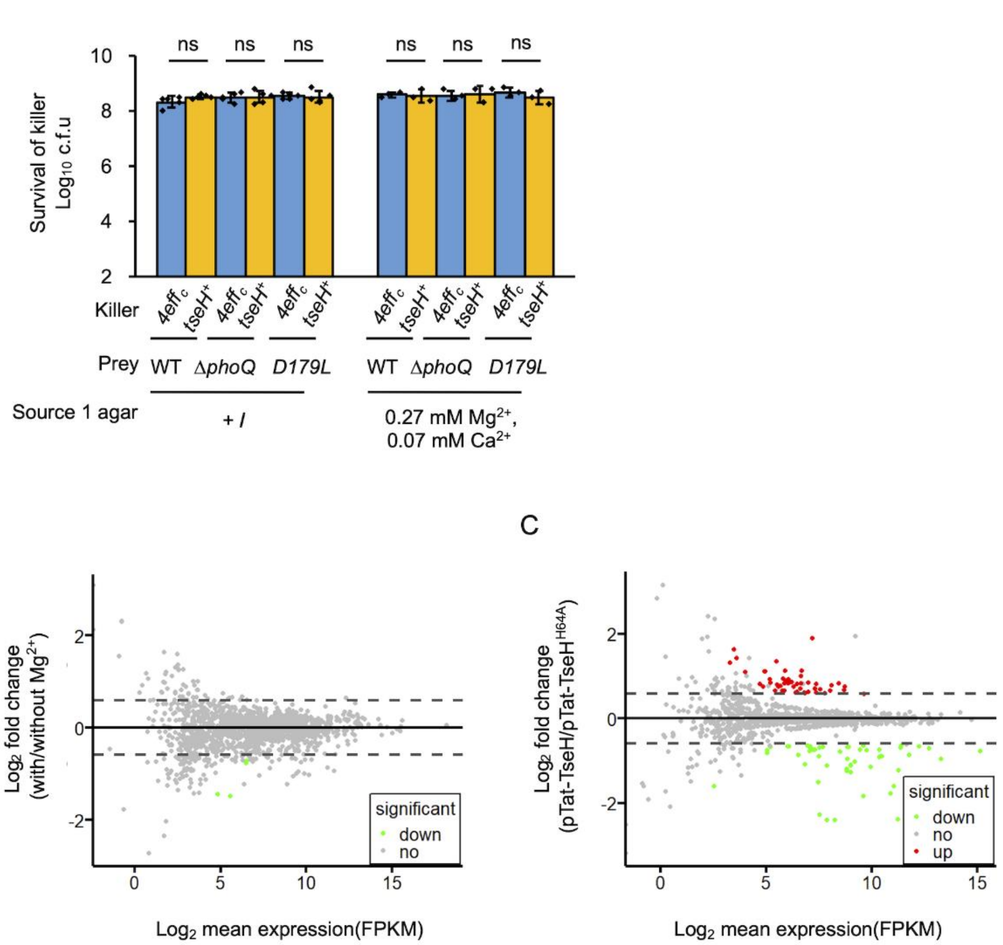


**Supplementary Figure 3. Stress response pathways protect *E. coli* against TseH. A**, Survival of killers after competition showed equal survival, relating to **Figure 3A**. Killer survival was selected by the streptomycin resistant on the genome. Error bars indicate the mean ± standard deviation of at least three biological replicates. Statistical significance was calculated using a two-tailed Student's *t*-test. ns, not significant. **B** and **C**, M-A plot of transcriptome results in *E. coli* samples with or without 0.3 mM $Mg^{2+}$(**B**) and *E. coli* samples ectopically expressing Tat-TseH and Tat-TseH$^{H64A}$(**C**). Up and down-regulated genes are indicated in red and green respectively. Genes with no significant change are indicated in gray.

Supplementary Figure 4：

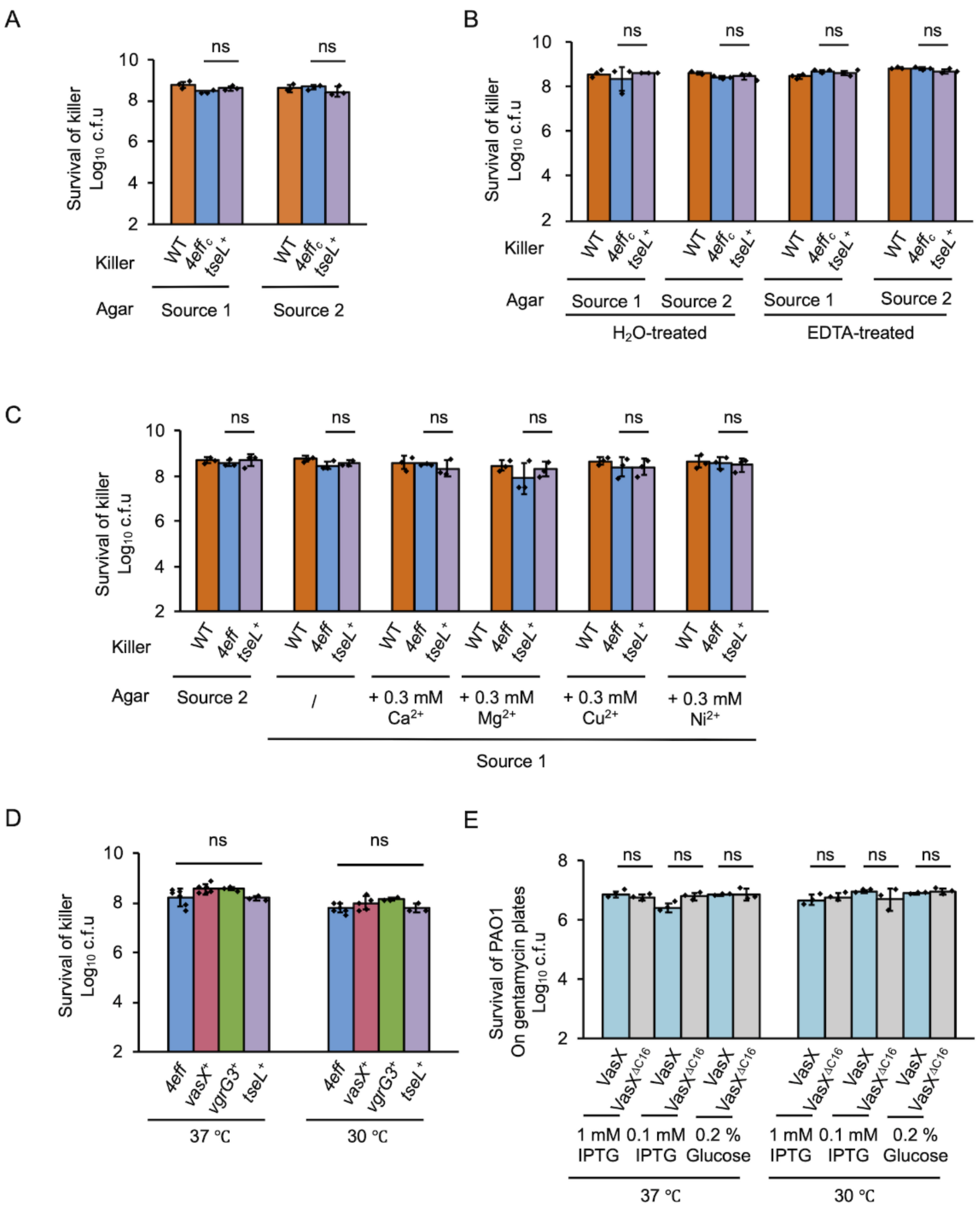


**Supplementary Figure 4. Effects of cations and temperature on effector-mediated killing.** **A-D**, survival of killer *V. cholerae* strains after competition showed equal survival, relating to **Figure 4A-4D**, respectively. Competition was performed on LB source-1-agar or source-2-agar plates (**A**), on LB EDTA-treated agar plates (**B**), on LB source-1-agar plates supplemented 0.3 mM $Ca^{2+}$, $Mg^{2+}$, $Cu^{2+}$ or $Ni^{2+}$ (**C**), and at 30 ℃ and 37 ℃ (**D**). For **A** to **D**, Killer strains are indicated at the bottom of each panel. WT, wild type; $4eff_C$, the 4-antibacterial-effector-inactive mutant; $tseL^+$, the TseL-active only mutant. $vasX^+$, the VasX-active only mutant. $vgrG3^+$, the VgrG3-active only mutant. Survival of killer strains was selected by the streptomycin resistant on the genome. **E**, Toxicity assay of *P. aeruginosa* PAO1 strains ectopically expressing Tat-VasX and its colicin-inactivated deletion mutant Tat-VasX$^{\Delta C16}$ on plates with gentamycin only at 30 ℃ or 37 ℃. For **A-E**, Error bars indicate the mean ± standard deviation of three biological replicates. Statistical significance was calculated using a two-tailed Student's *t*-test for two groups comparison or one-way ANOVA test for more than two groups comparison. ns, not significant;

Supplementary Figure 5：

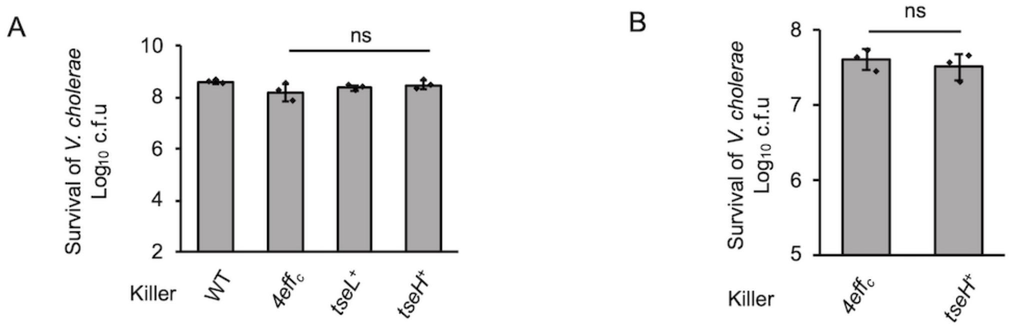


**Supplementary Figure 5. Survival of killers after competition on LB raw agar plates, relating to Figure 5. A,** Survival of *V. cholerae* strains after competition with *E. coli*. **B**, Survival of *V. cholerae* strains after competition with the Δ*tsiH* mutant. For **A** and **B,** WT, wild type; $4eff_c$, the 4-antibacterial-effector-inactive mutant; $tseL^+$, the TseL-active only mutant; $tseH^+$, the TseH-active only mutant. Survival of killer strains was selected by the streptomycin resistant on the genome. Error bars indicate the mean ± standard deviation of three biological replicates. Statistical significance was calculated using a two-tailed Student's *t*-test. ns, not significant.

**Supplementary Table 1**. Determination of element content in agar powder by XRF

| Agar (Source 1) | | Agar (Source 2) | |
|---|---|---|---|
| element | Content (%) | element | content (%) |
| C | 95.2992 | C | 97.8896 |
| Na | 1.9997 | S | 0.7530 |
| S | 0.9985 | Ca | 0.6196 |
| Cl | 0.8280 | Na | 0.3374 |
| P | 0.4775 | Mg | 0.1867 |
| K | 0.1256 | Si | 0.0549 |
| Si | 0.0690 | Cl | 0.0384 |
| I | 0.0447 | K | 0.0383 |
| Ca | 0.0381 | Ni | 0.0253 |
| Cu | 0.0321 | Cu | 0.0212 |
| Br | 0.0314 | Al | 0.0157 |
| Fe | 0.0267 | Fe | 0.0127 |
| Ni | 0.0195 | Sr | 0.0040 |
| Al | 0.0100 | P | 0.0030 |

**Supplementary Table 2.** Strains and plasmids used in this study.

| Strain | Genotype | Description | Source |
|---|---|---|---|
| *V. cholerae* V52 rhh | Parental | Deletion in *rtxA hlyA hapA*, parental strain | [1] |
| | Δ*vasK* | T6SS null, in-frame deletion of VCA0120 | [1] |
| | *vipA-mCherry2* | C-terminal chromosomal fusion of mCherry2 to VipA | [2] |
| | *vipA-mCherry2, tseL$^{D425A}$/ vgrG3$^{D842A}$/ vasX$^{\Delta C16}$/ tseH$^{H64A}$* | the *4eff$_c$* mutant, all four antibacterial effector inactive mutant | [3] |
| | *vipA-mCherry2, tseL$^{D425A}$/ vgrG3$^{D842A}$/ vasX$^{\Delta C16}$* | the *tseH$^+$* mutant, triple antibacterial effector inactive mutant in *tseL*, *vgrG3* and *vasX*, only TseH active | [4] |
| | *vipA-mCherry2, tseL$^{D425A}$/ vgrG3$^{D842A}$/ tseH$^{H64A}$* | the *VasX$^+$* mutant, triple antibacterial effector mutation in *tseL*, *vgrG3* and *tseH*, only VasX active | [5] |
| | *vipA-mCherry2, vgrG3$^{D842A}$/ vasX$^{\Delta C16}$/ tseH$^{H64A}$* | the *tseL$^+$* mutant, triple antibacterial effector mutation in *vgrG3*, *vasX* and *tseH*, only TseL active | [5] |
| | *vipA-mCherry2, tseL$^{D425A}$ / vasX$^{\Delta C16}$/ tseH$^{H64A}$* | the *vgrG3$^+$* mutant, triple antibacterial effector mutation in *tseL*, *vasX* and *tseH*, only VgrG3 active | [5] |
| | *vipA-mCherry2*, Δ*paar2-tseH-tsiH* | In-frame deletion of VCA0284-0286 | (this study) |
| | *vipA-mCherry2, tseL$^{D425A}$/ vgrG3$^{D842A}$/ vasX$^{\Delta C16}$*, Δ*paar2-tseH-tsiH* | Triple effector inactive and in-frame deletion of VCA0284-0286 | (this study) |
| *E. coli* | | | |
| pir1 | F-, Δ*lac169, rpoS(Am), robA1, creC510, hsdR514, endA,recA1, uidA(*Δ*MluI)::pir-116* | Strain used for cloning | Invitrogen |
| WM6026 | *lacIq , rrnB3, DlacZ4787, hsdR514, DaraBAD567, DrhaBAD568, rph-1, attl::pAE12(DoriR6K-cat::Frt5), DendA::Frt, uidA(DMluI)::pir, attHK::pJK1006D(oriR6K-cat::Frt5; trfA::Frt)* | Strain used for conjugation | Mekalanos Lab |
| DH5alpha | *F- Φ80lacZΔM15 Δ(lacZYA-argF) U169 recA1 endA1 hsdR17 phoA supE44 thi-1 gyrA96 relA1 λ-* | Strain used for cloning | Invitrogen |
| T-fast | *F' proA+ B+ lacIq* Δ*lacZM15 / fhuA2* Δ*(lac-proAB) glnV galK16 galE15 R(zgb-210::Tn10)TetR endA1 thi-1* Δ*(hsdS-mcrB)5* | Strain used for cloning | TIANGEN |

| | | | |
|---|---|---|---|
| MG1655 | *F-, lambda-, rph-1* | K-12 wild-type strain used as prey for T6SS killing | Schellhorn lab |
| MG1655 | *F-, lambda-, rph-1*, Δ*phoQ* | In-frame deletion of phoQ | (this study) |
| MG1655 | *F-, lambda-, rph-1, phoQ*$^{D179L}$ | Locked-on PhoQ mutant, constitutively pophosphorylates PhoP | (this study) |
| | Δ*rpoS* | | Lab stock |
| BW25113 | | Keio collection parental strain derived from MG1655 | [6] |
| *Aeromonas dhakensis* SSU | WT | Parental strain | [7] |
| | Δ*vasK* | T6SS null, in-frame deletion of *vasK* | [7] |
| *Pseudomonas aeruginosa* PAO1 | WT | | Mekalanos Lab |

### Plasmid

| | Plasmid | Description | Source |
|---|---|---|---|
| pDS132 | | Suicidal conjugation vector for all chromosomal allelic changes | [8] |
| | pDS132-MG1655-dphoQ | Suicidal vector to construct MG1655 chromosomal Δ*phoQ* | (this study) |
| | pDS132-MG1655-phoQ$^{D179L}$ | Suicidal vector to construct MG1655 chromosomal *phoQ*$^{D179L}$ | (this study) |
| | pDS132-MG1655-dphoP | Suicidal vector to construct MG1655 chromosomal Δ*phoP* | (this study) |
| | pDS132-V52-dvca0285-0286 | Suicidal vector to construct V52 chromosomal Δ*vca0285-0286* | (this study) |
| | pDS132-V52-dvca0284-0286 | Suicidal vector to construct V52 chromosomal Δ*vca0284-0286* | (this study) |
| pBAD24 | pBAD24Kan | Arabinose inducible expression plasmid, kanamycin resistance | Lab stock |
| | pBAD24Kan-V5 | Arabinose inducible expression plasmid, kanamycin resistance, C-terminal V5 tag | Lab stock |

| | | | |
|---|---|---|---|
| | pBAD24Kan-Tat-V5 | Arabinose inducible expression plasmid, kanamycin resistance, N-terminal Tat signal, C-terminal V5 tag | Lab stock |
| | pBAD24kan-MG1655-rcsA-V5 | Arabinose inducible expression of RcsA | (this study) |
| | pBAD24kan-MG1655-baeR-V5 | Arabinose inducible expression of BaeR | (this study) |
| | pBAD24kan-MG1655-spy-V5 | Arabinose inducible expression of Spy | (this study) |
| | pBAD24Kan-MG1655-clpB-V5 | Arabinose inducible expression of ClpB | (this study) |
| | pBAD24Kan-MG1655-yobF-V5 | Arabinose inducible expression of YobF | (this study) |
| | pBAD24Kan-MG1655-cspC-V5 | Arabinose inducible expression of CspC | (this study) |
| | pBAD24Kan-MG1655-phoQD179L-V5 | Arabinose inducible expression of $PhoQ^{D179L}$ | (this study) |
| | pBAD24kan-MG1655-phoQ-V5 | Arabinose inducible expression of PhoQ | (this study) |
| | pBAD24Kan-TseH-V5 | Arabinose inducible expression of TseH | (this study) |
| | pBAD24Kan-$TseH^{H64A}$-V5 | Arabinose inducible expression of $TseH^{H64A}$ | (this study) |
| | pBAD24Kan-Tat-TseH-V5 | Arabinose inducible expression of TseH with Tat signal | (this study) |
| | pBAD24Kan-Tat-$TseH^{H64A}$-V5 | Arabinose inducible expression of $TseH^{H64A}$ with Tat signal | (this study) |
| pBAD18 | pBAD18Cm | Arabinose inducible expression plasmid, choloramphenicol resistance | Lab stock |
| | pBAD18Kan | Arabinose inducible expression plasmid, kanamycin resistance | Lab stock |
| pPSV37 | | IPTG inducible expression plasmid, gentamycin resistant | Lab stock |
| | pPSV37-Tat | IPTG inducible expression plasmid, gentamycin resistant, N-terminal Tat signal | Lab stock |
| | pPSV37-Tat-VasX-FLAG | IPTG inducible expression of VasX in the periplasm | [4] |
| | pPSV37-Tat-$VasX^{\Delta C16}$-FLAG | IPTG inducible expression of $VasX^{\Delta C16}$ in the periplasm | [4] |
